# Federated deep learning enables cancer subtyping by proteomics

**DOI:** 10.1101/2024.10.16.618763

**Authors:** Zhaoxiang Cai, Emma L Boys, Zainab Noor, Adel Aref, Dylan Xavier, Natasha Lucas, Steven G Williams, Jennifer M S Koh, Rebecca C Poulos, Yangxiu Wu, Michael Dausmann, Karen L MacKenzie, Adriana Aguilar-Mahecha, Carolina Armengol, Maria M Barranco, Mark Basik, Elise D Bowman, Roderick J Clifton-Bligh, Elizabeth A Connolly, Wendy A Cooper, Bhavik Dalal, Anna DeFazio, Martin Filipits, Peter J Flynn, J Dinny Graham, Jacob George, Anthony J Gill, Michael Gnant, Rosemary Habib, Curtis C Harris, Kate Harvey, Lisa G Horvath, Christopher Jackson, Maija R J Kohonen-Corish, Elgene Lim, Jia Liu, Georgina Long, Reginald V Lord, Graham J Mann, Geoffrey W McCaughan, Lucy Morgan, Leigh C Murphy, Sumanth Nagabushan, Adnan M Nagrial, Jordi Navinés, Benedict J Panizza, Jaswinder S Samra, Richard A Scolyer, Ioannis Souglakos, Alexander Swarbrick, David M Thomas, Rosemary L Balleine, Peter G Hains, Phillip J Robinson, Qing Zhong, Roger R Reddel

**Author notes:** These authors contributed equally (Zhaoxiang Cai, Emma L Boys, Zainab Noor, Adel Aref).

## Abstract

Artificial intelligence applications in biomedicine face major challenges from data privacy requirements. To address this issue for clinically annotated tissue proteomic data, we developed a Federated Deep Learning (FDL) approach (ProCanFDL), training local models on simulated sites containing data from a pan-cancer cohort (n=1,260) and 29 cohorts held behind private firewalls (n=6,265), representing 19,930 replicate data-independent acquisition mass spectrometry (DIA-MS) runs. Local parameter updates were aggregated to build the global model, achieving a 43% performance gain on the hold-out test set (n=625) in 14 cancer subtyping tasks compared to local models, and matching centralized model performance. The approach’s generalizability was demonstrated by retraining the global model with data from two external DIA-MS cohorts (n=55) and eight acquired by tandem mass tag (TMT) proteomics (n=832). ProCanFDL presents a solution for internationally collaborative machine learning initiatives using proteomic data, e.g., for discovering predictive biomarkers or treatment targets, while maintaining data privacy.

**Statement of Significance:** A federated deep learning approach applied to human proteomic data, acquired using two distinct proteomic technologies from 40 tumor cohorts from eight countries, enabled accurate cancer histopathological subtyping while preserving data privacy. This approach will enable privacy-compliant development of large-scale proteomic AI models, including foundation models, across institutions globally.

## Introduction

Artificial intelligence (AI) applications, driven by a wealth of online data, have gained traction as tools to enhance efficiency, convenience, and innovation across multiple sectors. The use of these applications in commercial products, ranging from personalized recommendations in streaming services to generative AI tools such as ChatGPT has led to widespread uptake of these technologies and ongoing discussion about their appropriate use and regulation (1). Within the biomedical domain, numerous applications of AI are undergoing rapid development, including diagnostic prediction tools, interpretation of radiological and histopathological images, and methods of drug discovery (2). Although such tools offer promise, with efficiency gains and the potential for novel insights beyond those that can be achieved by traditional research studies, to date, large-scale AI modeling in most biomedical fields continues to be hindered by several substantive challenges (3).

These challenges include the privacy of data, especially personal clinical records, data ownership and governance, human research ethics, and intellectual property concerns (4). In transnational studies, compliance with laws and regulations that impose stringent standards on the collection, storage, sharing and use of biomedical data may be complicated by differing requirements in the relevant jurisdictions (5,6). Consequently, data sharing among collaborators within international consortia can be infeasible, hindering the assembly of heterogeneous and globally representative large-scale datasets, and presenting a significant barrier to the development of practical and relevant AI tools in the biomedical field. This contrasts sharply with commercial AI products based on ubiquitous information regarded as non-sensitive. However, for biomedical applications such as cancer research there is an urgent need to utilize all sources of good-quality data for purposes that include expediting the discovery of drug targets and predictive biomarkers, and avoid duplication of resources and therefore wastage due to data siloing and inaccessibility. To address these challenges, innovative solutions that balance the need for data accessibility and protection of clinical data are crucial.

Federated learning (FL) offers a promising solution to some of these challenges (7). As a distributed learning framework, FL permits local training of sensitive data at participating sites, with only the local model updates being shared with a central server to create a global model. This approach ensures that local data remains protected and securely stored behind firewalls. The ability to protect confidential data and combine diverse and geographically distinct datasets has outstanding potential for the development of large-scale generative AI tools with utility in both biomedical research and healthcare settings (8–10).

Although genomic and transcriptomic studies have greatly advanced our understanding of cancer, proteomic data will play a crucial role in answering many unresolved questions regarding the molecular mechanisms of cancer (11–13) and in identifying predictive markers (13). However, as the scale of human proteomics research increases, so do the challenges related to data privacy. FL provides a promising solution to address this, but so far has been applied only to non-human proteomic data (14). A potential high-impact application of FL in proteomics would be to develop a federated global model for international proteomic consortia, as will be attempted by π-Hub (the Proteomic Navigator of the Human Body) (15).

This study addresses these gaps by developing a federated deep learning (FDL)-based framework, ProCanFDL, for the analysis of proteomic data. The dataset, referred to here for brevity (and to distinguish it from external datasets) as the ProCan Compendium, includes 7,525 human biospecimens from 30 cohorts that were preserved and stored either by freezing or formalin fixation and paraffin embedding in pathology laboratories in multiple countries. There were sufficient samples to train the FDL proteomic model to recognise 14 cancer histopathological subtypes; its accuracy, tested on a hold-out test set, consistently outperformed individual local models, and were on par with the centralized model. The robustness of ProCanFDL was further validated using ten external proteomic datasets, eight of which were generated by a different MS technology, covering two additional cancer subtypes, to train a global model that can accurately recognise 16 histopathological subtypes. These findings highlight the potential of FDL to advance global clinical proteomics research by enabling secure, integrative data analysis across institutions and jurisdictions.

## Results

### ProCan Compendium and Landscape Analysis

We first compiled the ProCan Compendium, quantifying proteomes from 7,525 tissue samples, including 5,982 tumors, 1,512 tumor-adjacent normal samples and 30 benign samples from 4,956 individual patients (**Supplementary Table 1**). The data were generated in collaborative research projects involving 20 research groups across seven countries (**Figure 1a**) who provided biospecimens, stored either fresh frozen (FF) or formalin-fixed and paraffin-embedded (FFPE), and the associated clinical data. Utilizing a high-throughput workflow, 19,930 Data-Independent Acquisition Mass Spectrometry (DIA-MS) runs were used to obtain replicate proteomic data from the 7,525 samples (11,16–18). Raw DIA-MS data were processed and normalized using DIA-NN with a DIA-NN-generated spectral library, quantifying a total of 9,102 proteins. The number of proteins quantified per sample, grouped by tissue of origin, cancer type, and cancer subtype, is presented in **Supplementary** Figure 1. These samples encompassed 31 tissues of origin, 29 cancer histopathology types, and over 65 cancer subtypes, distributed across 30 cohorts (**Figure 1b, c, Methods**). High correlations between replicates of individual samples were observed, with a sample-wise median Pearson’s correlation coefficient (Pearson’s r) of 0.96, and moderate correlations between samples of the same cancer and tissue of origin (0.84 and 0.81, respectively). Correlations between unmatched samples from the same instrument were equivalent to those of random sample pairings (median Pearson’s r = 0.75), indicating that there were no instrument-specific batch effects (**Figure 1d**).

**Figure 1.**
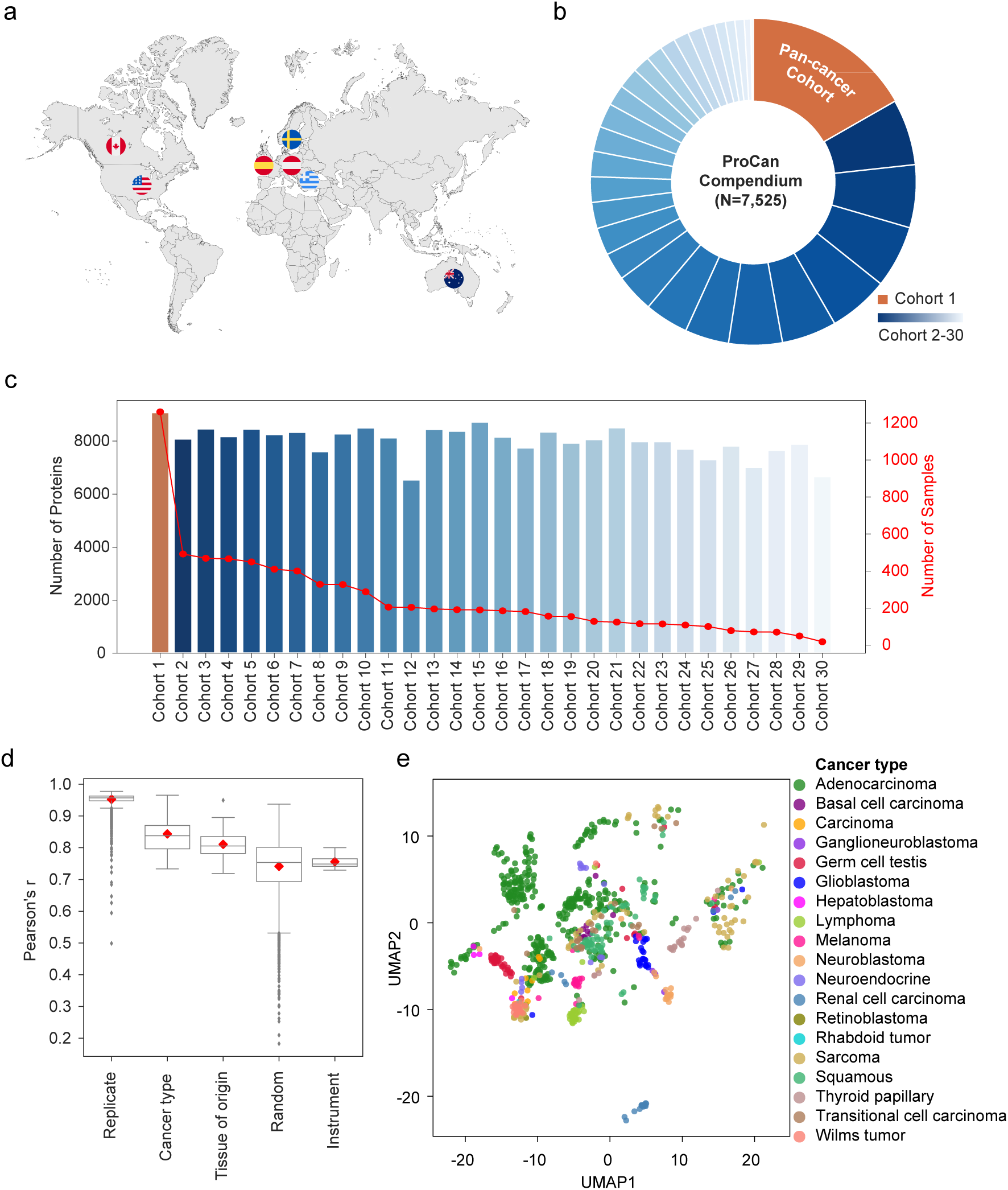
Overview of ProCan Compendium. **a**) The data were assembled from studies involving 20 collaborating cancer research groups across seven countries. **b**) The circular bar plot shows the sample sizes of the 30 cohorts, with the largest being the pan-cancer cohort (Cohort 1). **c**) Bar plot showing the number of quantified proteins; the red dot indicates the number of samples. **d**) Box plot of mean Pearson’s r for replicates, cancer types, tissue of origin, random sets of 10,000 samples, and different MS instruments. Box-and-whisker plots display 1.5 x interquartile ranges, with centers indicating medians and red diamonds representing mean values. **e**) UMAP with samples colored by cancer types.

In the ProCan Compendium, Cohort 1 serves as the baseline pan-cancer cohort and its raw data, and the corresponding spectral library are made publicly available alongside this study as a resource for researchers in the field of cancer proteomics (**Data Availability**). This cohort was acquired from the Victorian Cancer Biobank, the Gynaecological Oncology Biobank (GynBiobank) at Westmead Hospital and the Children’s Medical Research Institute Legacy sample set and consists of 766 tumor samples from 638 patients. Similarly to the ProCan Compendium overall, high correlations were observed between replicates of individual samples across all cancer types in Cohort 1 (**Supplementary** Figure 2a). Protein intensities were visualized in tumor samples using Uniform Manifold Approximation and Projection (UMAP), revealing distinct clusters for several cancer types, including adenocarcinoma, lymphoma, and melanoma, with proximate clustering observed for groups of related cancer types such as neuroblastoma and ganglioneuroblastoma, indicating the robustness of this pan-cancer dataset (**Figure 1e**). Analysis of cancer subtype-enriched proteins showed that Cohort 1 exhibited a pattern consistent with our previous study in cancer cell lines (13), with neuroblastoma showing the highest number of enriched proteins and quantification rate, and lymphoma the second highest (**Supplementary** Figure 2b,c). Cohorts 2–30 comprise 29 single-cancer cohorts, with a total of 5,217 tumor samples from 4,318 patients encompassing 42 cancer subtypes, which will be included in separate publications.

### ProCanFDL Overview

The traditional method of machine learning is based on local learning (**Figure 2a**), where individual research groups independently train models on the data available to them. This approach preserves jurisdictional data control but limits the ability to generalize findings across diverse datasets. Centralized learning (**Figure 2b**) improves predictive performance by aggregating data from multiple sites into a centralized model; however, it necessitates sharing sensitive data, with subsequent privacy concerns. FL (**Figure 2c**) represents an evolution of these methodologies by enabling the training of a global model across decentralized data sources, updating both local and global model weights without the need to transfer raw data, thus preserving data privacy. FDL then specifically refers to the implementation of deep learning techniques within this distributed setup.

**Figure 2.**
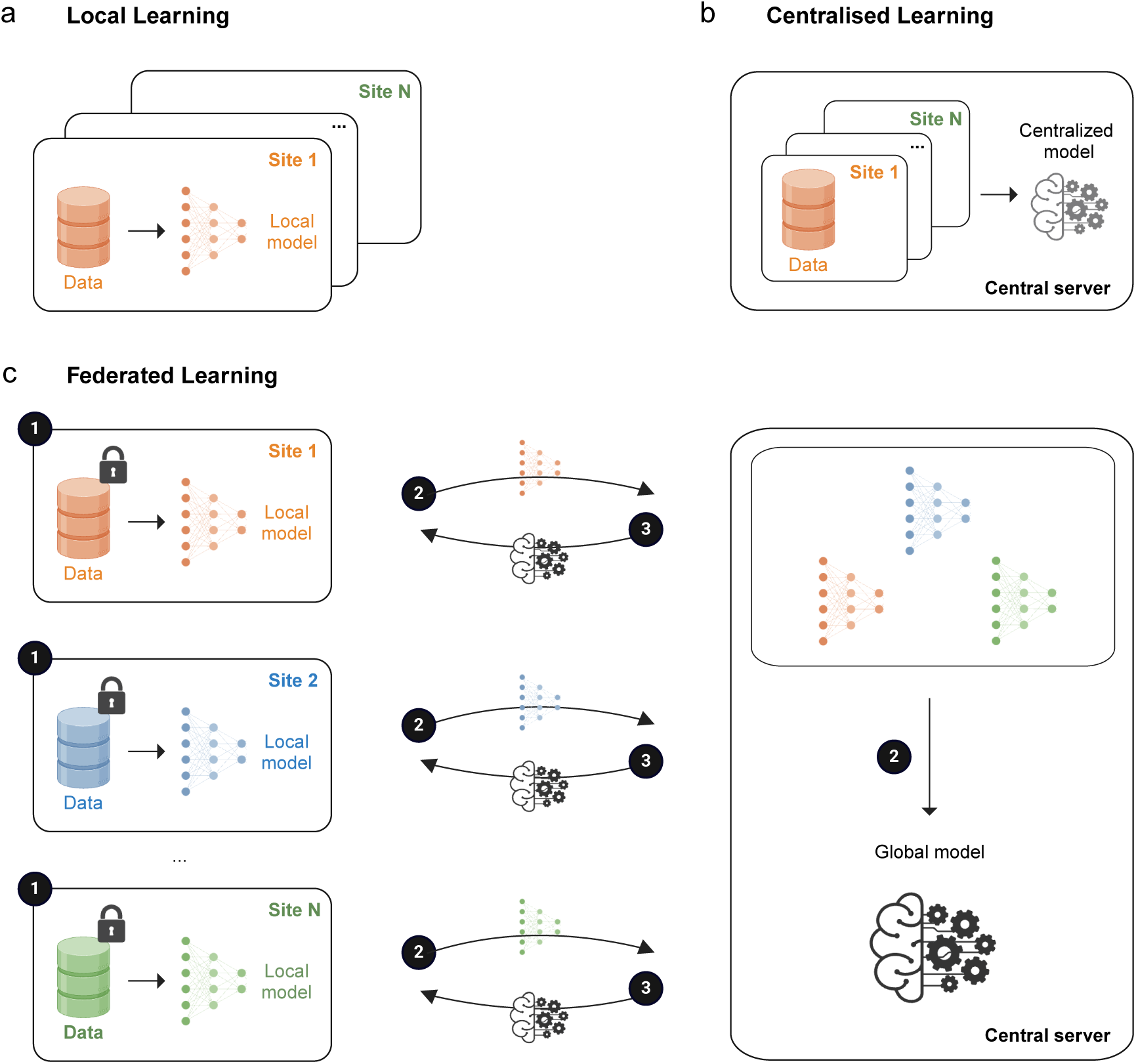
Local learning, centralized learning and federated learning. **a)** Local learning refers to training machine learning models on local sites without sharing data with other sites or a central server. **b)** Centralized learning involves collecting data from multiple local sites and aggregating data on a central server where a centralized machine learning model is trained. **c)** In federated learning, local models are trained on decentralized sites, each residing behind their respective firewalls. Only the model parameters, and not the raw data, are shared with a central server. These model parameters are then aggregated to form a global model. The global model is sent back to all local sites for the next round of training. The numbers 1 to 3 correspond to the first three algorithmic steps in ProCanFDL.

The ProCanFDL framework employs a four-step FDL approach while maintaining data privacy, enabling collaborative research in an international consortium (**Figure 2c, Methods**). In Step 1 (Initialization and local training), a global model is initialized with random weights and distributed to all participating local sites. Then, a local instance of a deep learning model is trained on its private proteomic data at each participating site. These models are trained independently, without sharing raw data across sites. In Step 2 (Global model aggregation), the trained model parameters are securely transferred to a central server, which aggregates these updates using a federated averaging algorithm. This process creates a global model reflecting the pooled knowledge from all local datasets, without the need for the server to access raw data. In Step 3 (Global model update), the newly aggregated global model is distributed back to all participating sites, where it serves as the starting point for the next round of local training. Finally, in Step 4 (Iteration and convergence), this process (Steps 1-3) is repeated iteratively, with each cycle refining the global model until it converges. The resulting model becomes increasingly accurate and representative of the combined datasets, encapsulating the collective knowledge.

### ProCanFDL on ProCan Compendium

To evaluate and benchmark ProCanFDL, we used proteomic data from the ProCan Compendium as input, training local, centralized, and ProCanFDL global models to assign each sample to its correct cancer subtype. As a proof of concept, we focused on 14 cancer subtypes, for each of which at least five samples were available in Cohort 1 and at least 20 samples across Cohorts 2-30. Additionally, only samples with replicate correlations greater than 0.9 were included in the analysis to ensure data quality and consistency. This filtering step resulted in a final subset of 4,558 samples, which was used in the subsequent analyses. No cohort-level normalization was performed. The input data for all machine learning models was organized in a data matrix format, where rows represent samples and columns correspond to protein abundances. We designated 10% of the patients from each of Cohorts 2-30 as the fixed hold-out test set *T*. The training set consisted of the remaining 90% of data from Cohorts 2-30 and all of the data from Cohort 1 (**Figure 3a, Methods**). The performance of the models for local, centralized, and ProCanFDL was evaluated using the same test set, *T*. Local, centralized and ProCanFDL models employed identical deep learning architecture and hyperparameters, optimized through the cross-validation process using Cohort 1 (**Figure 3b, Methods**).

**Figure 3.**
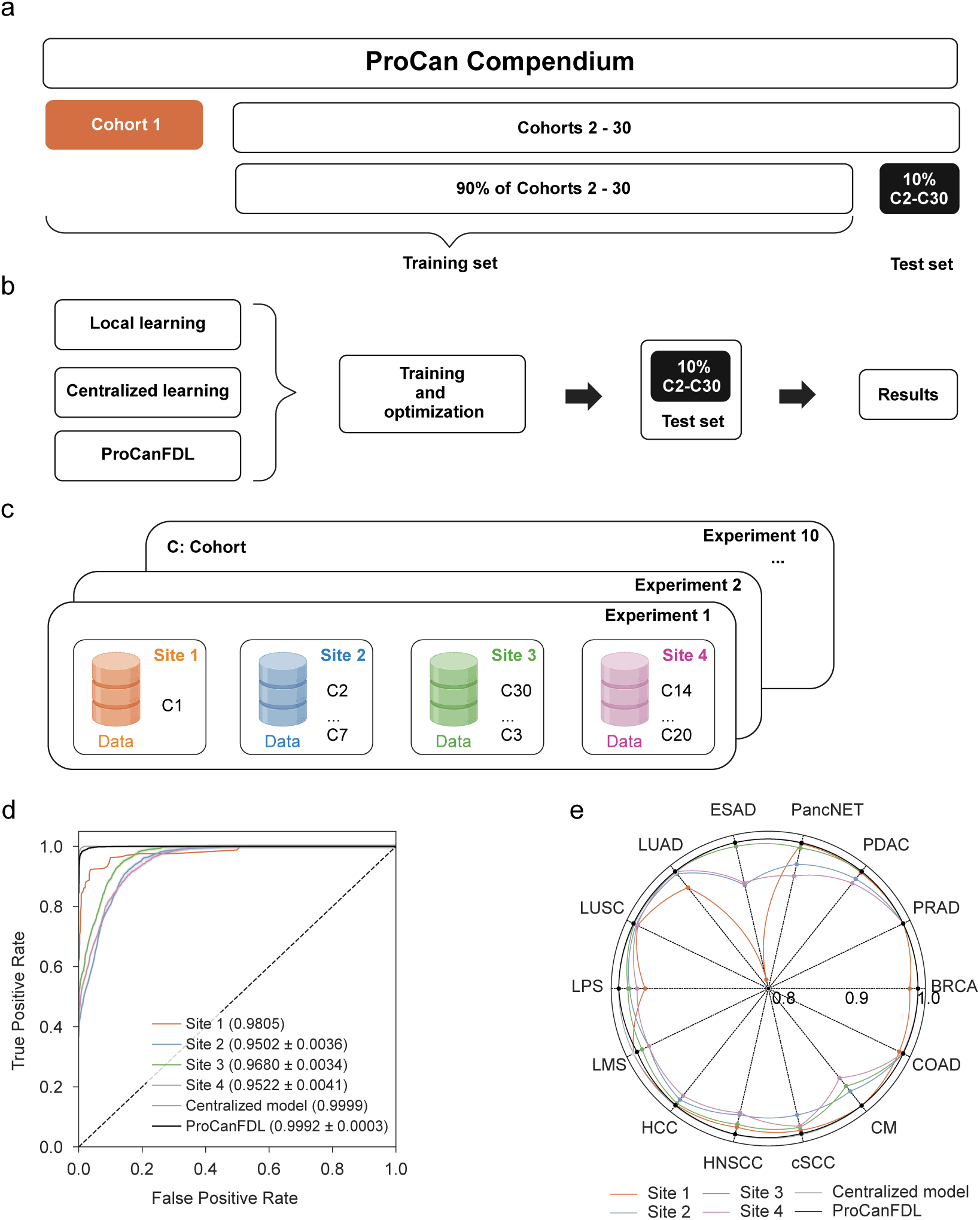
Experiment setup and model performance. **a)** The ProCan Compendium consists of Cohorts 1-30. Cohort 1, along with 90% of Cohorts 2–30, forms the training set, while the remaining 10% of Cohorts 2–30 constitutes the hold-out test set. **b)** The final local, centralized, and ProCanFDL models were evaluated on the same test set. **c)** Ten experiments for evaluating ProCanFDL. Site 1 always contains Cohort 1 (C1), while each site from Sites 2 to 4 contains a random subset of cohorts. **d)** Performance of each model was benchmarked by macro-averged AUROC for Sites 2-4 and ProCanFDL. The AUROCs were annotated as the mean values ± the half-width of the 95% confidence intervals (CI) estimated from ten experiments. **e)** AUROC of models across 14 cancer subtypes. The full names of the cancer subtypes can be found in **Methods**.

#### Site Simulation

We created four local sites to simulate a federated learning scenario where data from different institutions cannot be centrally combined due to privacy regulations. To obtain statistically robust and generalizable results, we ran the simulation ten times. In each iteration, Site 1 always included only data from Cohort 1, while the other sites received a randomly selected, non-repeating subset of cohorts from the remaining 29. This approach generated ten unbiased cohort distributions across Sites 2-4, ensuring that each site contained approximately ten cohorts, representing a meaningful fraction of the 14 cancer subtypes (**Figure 3c**).

#### Local Learning

We trained a local model for Cohort 1 data in Site 1. The performance of the final model achieved a macro-averaged area under the receiver operating characteristic curve (AUROC) of 0.9805 and an accuracy of 0.847 for classifying the 14 cancer subtypes (**Figure 3d, Supplementary** Figure 3a**, Methods**). We then evaluated the performance of the local models for Sites 2-4 by averaging the results from ten experiments for each site. Site 2 achieved a mean macro-averaged AUROC of 0.9502, and Sites 3 and 4 recorded values of 0.9680 and 0.9522, respectively (**Figure 3d**). These scores were lower than the mean macro-averaged AUROC of Site 1, primarily due to the absence of data encompassing all 14 cancer subtypes, which was only available at Site 1.

#### Centralized Learning

The centralized learning approach combines all data from the training set to train a single centralized model. The centralized model achieved a macro-averaged AUROC of 0.9999 (**Figure 3d**) and an accuracy of 0.990 (**Supplementary** Figure 3a), significantly outperforming the local models. These results demonstrate the expected benefits of data aggregation in improving predictive performance, but this approach requires the sharing of data between sites which may be prevented by local laws and regulations.

#### ProCanFDL

Using the four-step algorithmic procedure of ProCanFDL, we developed a global model for each experiment (**Methods**). The performances of the global models were averaged across the ten experiments. The ProCanFDL global model achieved a mean macro-averaged AUROC of 0.9992 (**Figure 3d**) and a mean accuracy of 0.965 (**Supplementary Figure 3a**). This represents a substantial improvement over the local models for Sites 1–4 and approximates the performance of the centralized model. The AUROC for each cancer subtype is detailed in **Figure 3e** and **Supplementary Table 2**. We next evaluated the number of true and false predictions across each class generated by the ProCanFDL global model with the highest macro-averaged AUROC from the ten experiments. The global model achieved 100% sensitivity (true positive rate) in classifying 10 out of 14 cancer subtypes, plus sensitivity exceeding 90% for lung adenocarcinoma (**Supplementary Figure 3b**), confirming the predictive power of the model.

Overall, these findings highlight the effectiveness of ProCanFDL in improving cancer subtyping performance. Notably, the federated approach not only surpasses the performance of local models but also delivers results comparable to the centralized model, while providing the critical advantage of preserving data privacy, potentially enabling large-scale machine learning across sites worldwide.

### Generalization and Integration

To rigorously assess the ProCanFDL model’s performance on unseen data and provide a clear indication of its generalizability beyond the ProCan Compendium dataset and the DIA-MS method, we included datasets generated in other laboratories. We selected datasets encompassing subsets of 65 cancer subtypes, generated using two distinct MS technologies (DIA-MS and Tandem Mass Tagging; TMT). For DIA-MS, we retrieved two cohorts from the PRoteomics IDEntifications (PRIDE) database, one consisting of 40 colorectal adenocarcinoma samples from Spain and another with 15 pancreatic ductal adenocarcinoma samples from South Africa (19,20). For TMT, we accessed data from the Clinical Proteome Tumor Analysis Consortium (CPTAC) in the USA, which included 832 tumor samples across eight cohorts (**Supplementary Table 3**) (21).

To ensure consistency across the different datasets, z-score normalization was applied to both the DIA-MS proteomic data (the ProCan Compendium and two external DIA-MS cohorts) and the eight TMT cohorts, separately. The two DIA-MS datasets were concatenated sample-wise into a single matrix before normalization, as were the eight TMT cohorts. The two normalized matrices were then concatenated sample-wise again to produce a final input matrix, containing a set of 3,837 proteins that were quantified in common in these cohorts.

We applied the train-test split across the external data (22). In this method, 90% of the external data, comprising the ten cohorts, was used to simulate additional local Sites 5 and 6, while the remaining 10% was held out and combined with the existing hold-out test set *T* to form a new, unbiased evaluation set *T′* **(Figure 4a)**. This test set *T′* was then used to evaluate and compare the generalizable performance of local, centralized, and ProCanFDL global models. The two external DIA-MS training cohorts were grouped as Site 5 and the eight TMT training cohorts from CPTAC as Site 6 **(Figure 4b)**. The inclusion of these external datasets allowed us to extend the analysis to two additional cancer subtypes, high-grade serous ovarian carcinoma and clear cell renal cell carcinoma, for which insufficient samples were available to meet the training cohort selection criterion of a minimum of 20 samples per cancer subtype in cohorts 2-30 of the ProCan Compendium. Thus, in the external validation analysis, the total number of cancer subtypes analyzed increased from 14 to 16, the total number of samples meeting the criteria for analysis in Sites 1-4 increased by 195 and the number of samples from Sites 1-4 included in the holdout test set increased by two.

**Figure 4.**
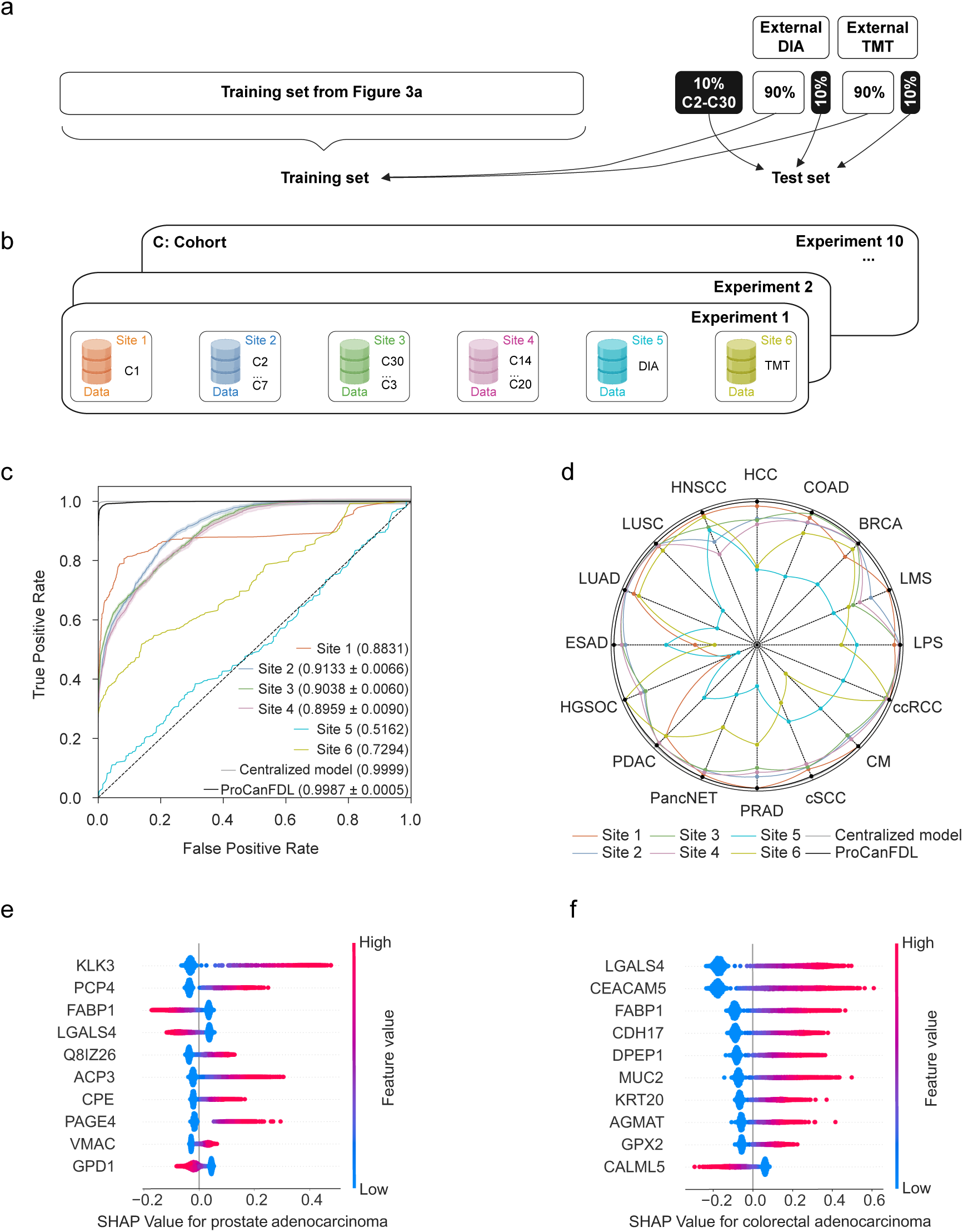
Generalization, integration and model interpretability. **a)** Illustration of the experimental design for inclusion of external datasets. The training data is expanded with 90% of each of the ten external datasets, while the test set is also expanded with 10% of the external data. **b)** Cohort allocations for the external datasets. The two external DIA-MS datasets are grouped in Site 5, while the eight external TMT datasets are grouped in Site 6. **c)** Performance of models was benchmarked by the AUROC with 95% CI estimated from the ten experiments. **d)** AUROC of models across 16 cancer subtypes. **e)** Beeswarm plot demonstrating the top 10 proteins contributing to the prediction of prostate adenocarcinoma. **f)** Top 10 proteins contributing to the prediction of colorectal adenocarcinoma.

The predictive performance of these local models was reflected by macro-averaged AUROCs of 0.8831 for Site 1, 0.5162 for Site 5, and 0.7294 for Site 6. Additionally, the mean macro-averaged AUROCs over ten experiments were 0.9133, 0.9038, and 0.8959 for Sites 2, 3, and 4, respectively (**Figure 4c**). Notably, local models from Sites 5 and 6 exhibited lower performance across different cancer subtypes compared to those from Sites 1-4, likely due to the limited coverage of the cancer subtypes. In contrast, the centralized model, trained on aggregated data from all six sites, exhibited significantly superior performance, achieving a macro-averaged AUROC of 0.9999 across all cancer subtypes (**Figure 4c**). By fully integrating both internal data (Sites 1-4) and external data (Sites 5 and 6), the centralized model captured a more comprehensive range of features and cancer subtypes, enhancing its predictive accuracy.

The ProCanFDL global model, trained in a federated manner across Sites 1-6, showed substantial improvements over the local models. The global model achieved a macro-averaged AUROC of 0.9987 (**Figure 4c**). The predictive performance of the global model was similar to that of the centralized model, while maintaining data privacy. Accuracy measures for local, centralized and ProCanFDL models are provided in **Supplementary Figure 4a**. The AUROC for each cancer subtype is detailed in **Figure 4d** and **Supplementary Table 4**. A confusion matrix was generated for the global model with the highest macro-averaged AUROC, providing a detailed assessment of true and false predictions for each subtype. The model achieved 100% sensitivity for 9 out of 16 cancer subtypes, with particularly high sensitivity rates for subtypes such as pancreatic ductal adenocarcinoma, hepatocellular carcinoma, and head and neck squamous cell carcinoma (**Supplementary Figure 4b**).

In summary, external validation of ProCanFDL, which also integrated datasets from two distinct MS platform types, showed that the global model consistently outperformed the local models. Importantly, the global model’s performance matched that of the centralized model.

### Model Interpretability

For potential downstream clinical application and interpretation, understanding the learned relationships and key discriminatory proteins in the ProCanFDL global model is important. By leveraging Shapley Additive Explanation (SHAP) values, we identified the top features contributing to the cancer subtype predictions made by the ProCanFDL global model, which achieved the highest AUROC when trained with internal data (**Supplementary Table 5)**. For example, proteins with utility at distinguishing between histological types of cancer were identified. Desmoglein 3 (DSG3), a marker of squamous differentiation, was identified within the top 10 SHAP values for both cutaneous and head and neck squamous cell cancers, with SHAP values for this marker contributing negatively towards the prediction of the other cancer subtypes (**Supplementary Figure 5a**). Similarly, markers suggesting epithelial differentiation, such as Anterior gradient 2, protein disulfide isomerase family member (AGR2) contributed positively towards the identification of breast, colorectal and pancreas adenocarcinomas, but negatively towards the identification of tumors arising from other cell types including pancreatic neuroendocrine tumors, sarcoma and melanoma (**Supplementary Figure 5b**). Finally, additional proteins were specific for identifying different tissues, including Kallikrein-related peptidase 3 (KLK3, Prostate-specific antigen) and Purkinje cell protein 4 (PCP4) for prostate adenocarcinoma (**Figure 4e**) as well as Galectin 4 (LGALS4) and Cadherin 17 (CDH17), which are known to be expressed in the intestinal epithelium, for colorectal adenocarcinoma (**Figure 4f**). Cytokeratin 20 (KRT20) featured within the top 10 SHAP values for predicting colorectal adenocarcinoma, and is known to have clinical utility for differentiating this cancer subtype from other subtypes of adenocarcinoma.

Over-representation analysis using the top 30 positive SHAP values for each cancer subtype revealed similar patterns with esophageal adenocarcinoma, pancreatic adenocarcinoma, leiomyosarcoma, colorectal adenocarcinoma, breast adenocarcinoma, liposarcoma and prostate adenocarcinoma demonstrating significant enrichment scores for their respective or closely related cancer subtypes (**Supplementary Table 5**). This was driven by proteins known to be enriched within specific cancer subtypes including breast cancer (Calmodulin like 5 (CALML5), Perlipin 1 (PLIN1) and Fatty acid binding protein 4 (FABP4)), colorectal cancer (LGALS4, Fatty acid binding protein 1 (FABP1)) and pancreatic adenocarcinoma (LGALS4, Peripherin (PRPH)). Of note, in the FDL model, one liposarcoma case was misclassified as breast carcinoma. The invasive lobular breast carcinoma pathway was significantly enriched in the top 30 proteins for liposarcoma due to the presence of proteins enriched in adipose tissue (Fatty acid binding protein 4, Perilipin 1 and Lipase E) in both cancer subtypes.

Next, we examined the protein features contributing positively to the prediction of each cancer subtype using the Hallmark gene set collection (23). For pancreatic adenocarcinoma, we noted proteins specific to pancreatic beta cells including Glucagon, Somatostatin and Pyruvate kinase L/R, as well as proteins related to up-regulation of KRAS signaling including Integrin subunit alpha 2, and several matrix metallopeptidases. For breast adenocarcinoma, we noted proteins related to estrogen response including several ETS transcription factors (**Supplementary Table 5**).

Collectively, the proteins highlighted by model interpretability analysis demonstrate that the FDL model predictions are plausibly related to known biological patterns that differ among cancer subtypes.

## Discussion

We have developed ProCanFDL, which enables accurate cancer subtyping using proteomic data derived from 7,525 FF and FPE human biospecimens. The framework leverages data from 20 cancer research groups across seven countries and processed in many different pathology laboratories, utilizing a privacy-preserving machine learning approach. Additionally, we demonstrated that ProCanFDL can integrate proteomic data generated via two different MS technologies.

The use of proteomic data in large-scale machine learning models has to date been limited by significant challenges, including the difficulties of integrating proteomic data from different platforms (24,25). Moreover, data centralization is difficult to achieve with sensitive patient data, especially from different jurisdictions (7). FL offers a potential solution to these issues by allowing collaborative training across multiple sites without the need to transfer raw data. The use of FL with patient data including chest X-rays and scans during the COVID-19 pandemic (10) illustrated the rapid gains that can be made by privacy-compliant data sharing.

Our model, which simulated FL by distributing data across four local sites behind private firewalls, demonstrated the feasibility of a real-world scenario whereby datasets are held across different institutions. By performing training independently at each local site and aggregating model updates centrally to form a global model, sensitive patient data can remain behind institutional firewalls. Notably, the model was enhanced by the incorporation generated by a distinct type of proteomic technology, TMT. Difficulties with the integration of data from different proteomic platforms have been a major issue in the aggregation of proteomic data. Therefore, ProCanFDL addresses multiple barriers to significantly scaling up proteomic machine learning analyses.

We used the macro-averaged AUROC as the primary model evaluation metric in this study. This method provides a more nuanced view of model performance compared to accuracy, especially in multi-class classification tasks where class imbalances may exist. By assigning equal weighting to each class, regardless of sample size, it ensures that performance is assessed fairly across all cancer subtypes, preventing larger classes from dominating and biasing the evaluation metrics.

As with previous FL studies (10), the ProCanFDL global model demonstrated a 43% improvement over local models, highlighting the benefits of aggregating model parameters from local sites to enhance sample diversity and representation of cancer subtypes. This collaborative approach is particularly advantageous for sites with limited samples or underrepresented subtypes. By incorporating data from multiple local sites, the global model could accurately predict subtypes that were not present in the local datasets, thereby increasing its robustness and generalizability. Additionally, the global model achieved comparable evaluation metrics to the centralized model, while offering the significant advantage of eliminating the need for data sharing between local sites. These attributes make ProCanFDL a practical and scalable solution for collaborative learning in proteomics, especially in situations where data privacy is paramount. This advantage is further underscored when considering the limitations of “regulatorily clean” datasets from some local sites, where patient selection criteria often result in cohorts that are not representative of broader patient populations.

Unlike most studies that use independent datasets with uniform distribution for external validation, we applied a train-test split across external datasets to address the inherently non-uniform data distributions obtained from different proteomic platforms and to ensure model generalization and robustness. This approach allows the model to adapt to variations in data while maintaining privacy.

By model interpretability analysis, we identified several important biological relationships including enrichment of proteins specific for cancer subtypes and relevant biological pathway information. This suggests that in addition to achieving high accuracy, the model can also capture meaningful biological signals. Such model explainability is vital for the acceptance of such technologies and potential translation into clinical practice. We anticipate that application of ProCanFDL will facilitate our understanding of cancer biology and treatment targets, and enable the use of proteomic data as an adjunct to histopathology in challenging diagnostic situations.

Although we demonstrated the utility of ProCanFDL, several limitations of the study provide scope for improvement and extension. One area for future research is the application of this FDL framework to more complex multi-institutional setups, where even greater variations in data types, collection methods, and processing protocols could introduce additional data harmonization challenges. Further, this study served as a proof-of-concept focused on cancer subtyping due to the availability of relevant annotations. Expanding its application to areas such as prognostic biomarker development and clinical outcome prediction will require datasets annotated with treatment outcomes, survival data and other detailed metadata at each participating site.

ProCanFDL enables the development of foundational models for proteomics. Unlike large language models such as ChatGPT or foundation models for digital pathology, which benefit from large public datasets or readily available imaging data, the development of large proteomic models faces significant challenges due to restricted access to clinically annotated datasets. ProCanFDL will enable the gathering and utilization of the data needed to train proteomic foundational models without compromising data privacy.

Overall, ProCanFDL represents a significant and tangible step toward applying federated learning to large-scale cancer proteomic datasets and generated by different MS technologies. By balancing the need for robust and accurate model performance with data privacy, it fosters a practical and scalable solution for proteomic data analysis and collaborative biomedical research. We anticipate that this will create new opportunities in cancer research, e.g., for discovery of novel treatment targets and predictive biomarkers, accelerate the clinical application of proteomic technologies, and extend to multi-omic data applications beyond proteomics.

## Methods

### Biospecimen and data collection

FF and FFPE samples were obtained from malignant samples (tumor and pre-malignant samples) and non-malignant tissues (benign tumors and tumor-adjacent normal samples). Ethics approval was obtained for use of all patient samples. Cohort 1 consisted of FF samples (n = 766 primary tumor samples and n = 494 tumor-adjacent normal samples) obtained from the Victorian Cancer Biobank (2019/ETH02039 (HREC/17/WMEAD/63), the Gynaecological Oncology Biobank (GynBiobank) at Westmead Hospital (2019/ETH02039 (HREC/17/WMEAD/63); 2019/ETH02043 (LNR/16/WMEAD/291)), and the Children’s Medical Research Institute Legacy sample set (2019/ETH05866 (LNR/17/SCHN/291). Cohorts 2-30 included both FF and FFPE samples obtained from the following sites: Gynaecological Oncology Biobank (GynBiobank) at Westmead Hospital, Australia (2019/ETH02039 (HREC/17/WMEAD/63); 2019/ETH02043 (LNR/16/WMEAD/291)), National Cancer Institute Biobank, United States of America (2019/ETH02039 (HREC/17/WMEAD/63); 2019/ETH02075 (LNR/17/WMEAD/249)), Westmead Institute for Medical Research, Australia (2019/ETH02039 (HREC/17/WMEAD/63); 2019/ETH10764 (LNR/19/WMEAD/39)), Institut Germans Trias i Pujol Research Institute, Badalona, Spain (2019/ETH06112 (HREC/17/SCHN/63)), PI-17-079), Royal Prince Alfred Hospital and Centenary Institute, Australia (2019/ETH02039 (HREC/17/WMEAD/63); 2021/ETH11460), Royal Prince Alfred Hospital and Woolcock Institute, Australia (2019/ETH02039 (HREC/17/WMEAD/63); 2020/ETH01304 (X20-0223)), St Vincent’s Hospital, Department of Surgery, Sydney, Australia and Institute of Clinical Sciences, Lund University Hospital, Sweden (2019/ETH02039 (HREC/17/WMEAD/63); 2021/ETH11590), Garvan Institute of Medical Research (APGI), Australia 2019/ETH02039 (HREC/17/WMEAD/63); X16-0293 (HREC/11/RPAH/329), Garvan Institute of Medical Research (2019/ETH02039 (HREC/17/WMEAD/63)), Melanoma Institute of Australia (2019/ETH02039 (HREC/17/WMEAD/63); X15-0454 (HREC/11/RPAH/444); X17-0312 (HREC/11/RPAH/32); X15-0311 (HREC/10/RPAH/530)), International Sarcoma Kindred Study (2019/ETH02039 (HREC/17/WMEAD/63); SVH 16/126; PMCC 09/11), University of Manitoba Tissue Biobank, Canada (2019/ETH02039 (HREC/17/WMEAD/63); HS14811 (H2001-083)), Australian Breast Cancer Tissue Bank (2019/ETH02039 (HREC/17/WMEAD/63); 2019/ETH02413 (LNR/16/WMEAD/93)), Nepean Research Biobank, Australia (2019/ETH02039 (HREC/17/WMEAD/63)), Princess Alexandra Hospital, Queensland Medical Labs, Mater Hospital, Queensland, Australia (2019/ETH02039 (HREC/17/WMEAD/63; PR/2022/QMS/8692 (HREC/03/QPAH/197), Institute of Cancer Research, Comprehensive Cancer Centre, Medical University of Vienna, Austria (2019/ETH02039 (HREC/17/WMEAD/63; 1312/2022), The University of Sydney, Royal North Shore Hospital, NSW Health Pathology, Australia (2019/ETH02039 (HREC/17/WMEAD/63; 2019/ETH08639 (HREC/16/HAWKE/105)), Jewish General Hospital Breast Cancer Biobank, Montreal, Canada (2019/ETH02039 (HREC/17/WMEAD/63); 2023-3377), Laboratory of Translational Oncology, Medical School, University of Crete, and Laboratory of Pathology, University Hospital of Heraklion, Greece (2019/ETH02039 (HREC/17/WMEAD/63); University of Crete ref 27/17.02.2020. University Hospital ref 9920).

Samples were sectioned as follows: 30-micron curls (at least one) for fresh frozen tissue and 10- to 20-micron curls (at least one) for FFPE tissue. A small proportion of samples underwent tissue punching (using a 1mm biopsy punch tool) or macrodissection to enhance the proportion of tumor content.

Specimens were annotated according to the provided histological cancer type and subtype diagnosis. For each sample, an adjacent hematoxylin and eosin slide was reviewed by a specialist pathologist to confirm that the section received in the proteomic laboratory was consistent with the diagnosis, and to evaluate the percentage of tumor content and necrosis, the extent of lymphocytic infiltration and the presence of additional tissue elements. Samples in Cohort 1 that were not consistent with the provided diagnosis (n = 84) and/or contained a significant presence of atypical or normal tissue elements (n = 116), had percentage tumor content <20% (n = 57) or percentage necrosis >80% (n = 14) were excluded from the FDL analyses.

### Sample preparation and mass spectrometric acquisition

All samples were prepared using the Heat ‘n Beat method and MS data were acquired from technical duplicate or triplicate MS runs (13).

### Spectral library generation

To generate the spectral library, 19,930 DIA-MS runs from the ProCan Compendium were collected in wiff file format and were processed using DIA-NN software. MS/MS spectra were referenced to the UniProt Human proteome. The spectral library containing 193,354 peptides including retention time peptides and peptides from commonly occurring microbial and viral proteins, and corresponding to a total of 15,306 proteins, was used to search the entire set of 19,930 sample and quality control runs to extract the DIA data.

### Data extraction

Raw DIA-MS data were processed using DIA-NN software, implementing retention time (RT) dependent normalization and the DIA-NN-generated spectral library. The input parameters are given below:

~~~
-report-lib-info --out step3-out.tsv --qvalue 0.01 --pg-level 1 --mass-acc- ms1 40 --mass-acc 40 --window 9 --int-removal 1 --matrices --temp . --smart- profiling --peak-center
~~~

Data were filtered to retain only precursors from proteotypic peptides with Global.Q.Value ≤ 0.01. Protein abundance was calculated using maxLFQ, with default parameters and implemented using the DIA-NN R Package (https://github.com/vdemichev/diann-rpackage). Data were then log2-transformed.

### Proteomic profiling

All data were processed using custom R/Python scripts. For the entire set of samples from the ProCan Compendium, the Pearson’s correlation coefficient was calculated using the corr function in Python package pandas (v2.0.3) to analyze the technical reproducibility of the data. In addition to the correlation among replicates from each sample, mean correlation among samples from different cancer types, tissue of origins, MS instruments and randomly selected samples was also calculated. Batch effects and clustering for these multi-cohort data were visualized using the UMAP dimensionality reduction tool. Moreover, the numbers of proteins quantified across tissue of origins and cancer types were assessed using the boxplots. The boxplots showed the range of proteins quantified in each tissue and cancer type along with median protein count in each class.

To further investigate the proteomic profiles of cancer subtypes from various origins, we selected cell-type enriched proteins using previously defined thresholds (13). Cell type-enriched proteins were defined as proteins quantified in at least 50% of samples from no more than one cancer subtype, and in ≦35% of samples from all other subtypes, considering only subtypes represented by at least 10 samples. For this, only tumor samples were used.

### Preprocessing and statistical analysis

For downstream analysis, the sample replicates were merged and a final protein matrix with only tumor samples was used. The protein matrix showed an average of 57% missingness per individual sample. Missing values were imputed with zero. No additional normalization or preprocessing was performed. In addition to filtering by replicate correlation, we also filtered samples in Cohort 1 to only include samples in the FDL analyses that met the following criteria: (1) consistent with the histopathological diagnosis, (2) adequate percentage of tumor content and (3) low percentage of necrosis (See **Methods, Biospecimen and Data Collection**).

### Preparation of training and test sets

The train-test split of 90% training and 10% testing was performed at the patient level, ensuring that multiple samples from the same patient were consistently assigned to the training set. For patients assigned to the test set, one random sample was selected to simulate real-world conditions.

### Hyperparameter tuning

Hyperparameter tuning was conducted via a 3-fold cross-validation on Cohort 1. Based on the cross-validation result, which helped identify the optimal architecture for model performance, the final architecture includes an input layer, a hidden layer, a ReLU activation function, a dropout layer with a probability of 0.2, and an output layer. The hyperparameters for the training process were set as follows: a learning rate of 1x10^-4^, a weight decay of 1x10^-4^, a hidden dimension size of 256, a batch size of 100, and a total of 200 epochs. The Adam optimizer was utilized to update the model parameters during training. Default settings were used for other hyperparameters that are not specifically mentioned above. These hyperparameters were then used for all models in local, centralized and ProCanFDL learning.

### ProCanFDL

ProCanFDL is a deep learning-based FL framework with a neural network architecture. FDL was conducted through iterative communication rounds between the central server and the participating local sites. The training procedure comprised the following four steps:

**Step 1 Initialization and Local Training**. A global model was first initialized with random weights and distributed to all participating local sites. This initial global model served as the starting point for the subsequent rounds of federated learning, ensuring that each site began the process with a common starting point. The model architecture was consistent across all sites, ensuring uniformity in training and subsequent aggregation. Each participating site locally trained its own instance of the deep learning model on its private proteomic data. PyTorch (v2.3.0) was utilized as the deep learning framework to implement and train the model. During this phase, model parameters (weights and biases) were optimized using the Adam optimizer. To prevent overfitting and enhance generalization, techniques such as early stopping and dropout were applied, as detailed in the model’s hyperparameter setup. The local models were trained independently, capturing unique proteomic signatures relevant to specific cancer types and tissues of origin. No raw data were shared between sites, ensuring data privacy.

**Step 2 Global Model Aggregation**. Following local training, the optimized model parameters, specifically the weights and biases, were securely transferred from each site to a central server. The server aggregated these updates using the federated averaging algorithm. The aggregation involved averaging the weights from all participating sites, resulting in a new global model that reflected the pooled knowledge from all local datasets, without the central server accessing any raw data. This step allowed the global model to capture the diversity of the proteomic data from all sites.

**Step 3 Global Model Update**. Once the aggregation was complete, the updated global model parameters were distributed back to all participating sites. Each site received the updated global model, which served as the starting point for the next round of local training. This iterative exchange allowed the model to progressively improve and adapt to the heterogeneous data across the sites.

**Step 4 Iteration and Convergence**. Steps 1 to 3 were repeated for a total of ten iterations. This fixed number of iterations allowed the model to progressively refine its performance by incorporating data from all local sites. After the ten iterations, the global model was evaluated on a hold-out test set to assess its generalizability across cancer subtypes. All global models in this study converged within ten iterations, but the number of iterations may need to be increased for other data and tasks.

A pseudocode for this four-step algorithm is described below:

~~~
// Initialization
Initialize global_model with random weights Distribute global_model to all local sites
// Iterative process with 10 iterations (Step 4)
for iteration in range(10):
// Step 1: Local Training
for each site in participating_sites:
// Train the local model using the given data and hyperparameters
local_model = train_model(global_model, site_data, hyperparameters)
end
// Optimize the model parameters using the Adam optimizer
local_weights = optimize(local_model, ’Adam’, hyperparameters)
// Step 2: Global Model Aggregation
// Send local model parameters (weights) to the central server
local_weights = send_to_server(local_model_parameters)
// Central server aggregates all local weights
global_weights = federated_average(local_weights)
end
// Step 3: Global Model Update
// Update global model with aggregated weights from all sites
global_model.update(global_weights)
// Distribute updated global model back to the local sites
distribute(global_model, participating_sites)
// Evaluation of the final global model on the test set evaluate(global_model, hold_out_test_set)
~~~

### Cancer subtypes for ProCanFDL

**BRCA** Breast carcinoma. **ccRCC** Clear cell renal cell carcinoma. **COAD** Colorectal adenocarcinoma. **CM** Cutaneous melanoma. **cSCC** Cutaneous squamous cell carcinoma. **ESAD** Esophageal adenocarcinoma. **HNSCC** Head and neck squamous carcinoma. **HCC** Hepatocellular carcinoma. **HGSOC** High-grade serous ovarian carcinoma. **LMS** Leiomyosarcoma. **LPS** Liposarcoma. **LUAD** Lung adenocarcinoma. **LUSC** Lung squamous carcinoma. **PancNET** Pancreatic neuroendocrine tumor. **PDAC** Pancreatic ductal adenocarcinoma. **PRAD** Prostate adenocarcinoma.

### Evaluation metrics

The performance of ProCanFDL was measured using the following two metrics. The first was the Area Under the Receiver Operating Characteristic Curve (AUROC), which is calculated for multi-class classification using the One-vs-Rest (OvR) approach. For each cancer subtype successively, that subtype is treated as the positive class while the remaining subtypes are considered to be negative, allowing for the calculation of per-class AUROC. To more comprehensively evaluate the model’s ability to discriminate between multi-class cancer subtypes, the macro-average AUROC is computed by averaging the AUROC scores across all classes without class-size weighting. This macro-average provides an overall measure of model performance across all classes, treating each class equally regardless of its prevalence in the dataset. The second metric is the multi-class Accuracy, which measures the proportion of correctly classified cancer subtypes among the total instances:

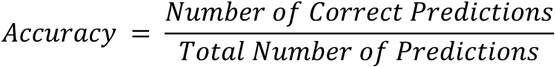

This provides a single value that summarizes the model’s performance across all cancer subtypes.

### Model interpretability analysis

Feature importance scores from the FDL model were calculated using Shapley Additive Explanation (SHAP) values with the Python package, SHAP (v0.45.1). Over-representation analysis using the top 30 proteins as indicated by positive SHAP values was performed using the DisGeNET database via Metascape (v3.5). For all over-representation analysis, the p-value cut-off was set as 0.05 and the input background gene set encompassed all proteins used for building the ProCanFDL model. Proteins with a positive SHAP value for each cancer subtype were also evaluated for potential biological relevance using the Hallmark gene set collection (23). In the beeswarm plots, features contributing positively to class prediction are shown on the right-hand side and features contributing negatively are shown on the left-hand side.

### Validation by external datasets

To provide consistency in the underlying model structure and training process, all local models, the centralized model and ProCanFDL global model were trained using the same model architecture and hyperparameter configuration as previously applied in local, centralized and federated learning. Z-score normalization was applied using the StandardScaler from scikit-learn (v1.4.2) to both the DIA proteomic data (19,20), and the eight TMT datasets from CPTAC (21) separately. Specifically, we first concatenated sample-wise all the DIA proteomic data (ProCan Compendium and two external DIA datasets) as one matrix, and then applied z-score normalization on this matrix. Similarly, the eight TMT cohorts were concatenated sample-wise into a single matrix, and z-score normalization was applied. Finally, the two normalized matrices were concatenated sample-wise to serve as the input for ProCanFDL.

## Data Availability

The raw DIA-MS data of Cohort 1 and the corresponding spectral library will be deposited to the Proteomics Identification Database (PRIDE). The two external DIA-MS datasets have the identifiers PXD019549 and PXD007810, respectively. Proteomics data for CPTAC datasets is available at Proteomic Data Commons (PDC). The PDC accession numbers for CPTAC datasets are: BRCA – S060; CCRCC – PDC000471; COAD – S045; HNSCC – PDC000221; LUAD – PDC000153; PDAC – PDC000270; LSCC – PDC000234; OV – S038.

## Acknowledgments

ProCan is supported by the Australian Cancer Research Foundation, Cancer Institute New South Wales (NSW) (2017/TPG001, REG171150), NSW Ministry of Health (CMP-01), The University of Sydney, Cancer Council NSW (IG 18-01), Ian Potter Foundation, the Medical Research Future Fund (MRFF-PD), National Health and Medical Research Council (NHMRC) of Australia European Union grant (GNT1170739, a companion grant to support the European Commission’s Horizon 2020 Program, H2020-SC1-DTH-2018-1, iPC - individualizedPaediatricCure), and National Breast Cancer Foundation (IIRS-18-164). The work at ProCan was done under the auspices of a Memorandum of Understanding between Children’s Medical Research Institute and the U.S. National Cancer Institute’s International Cancer Proteogenomics Consortium (ICPC), that encourages cooperation among institutions and nations in proteogenomic cancer research in which datasets are made available to the public. The Victorian Cancer Biobank through the Cancer Council Victoria as Lead Agency is supported by the Victorian Government through the Victorian Cancer Agency- a business unit of the Department of Health and Human Services. GVL is supported by an NHMRC Investigator Grant (2021/GNT2007839) and by the University of Sydney Medical Foundation. RCP and PJR are supported by NHMRC Fellowships (GNT1138536 and GNT1137064, respectively). RCP is supported by a Sydney Cancer Partners Translational Partners Fellowship with funding from a Cancer Institute NSW Capacity Building Grant (grant ID 2021/CBG0002). ZC is the recipient of a PhD Scholarship from Sydney Cancer Partners with funding from Cancer Institute NSW (2021/CBG0002). RAS is supported by a National Health and Medical Research Council of Australia (NHMRC) Investigator Grant (GNT2018514). Research conducted at the Westmead Institute for Medical Research (WIMR) was supported by Tour de Cure and the Cancer Institute NSW (CINSW) through the Sydney West Translational Cancer Research Centre (SW-TCRC, 15/TRC/1-01). Tissues were received from the Australian Breast Cancer Tissue Bank (ABCTB) which is generously supported by the NHMRC, the Cancer Institute NSW and the NBCF. The Gynaecological Oncology Biobank at Westmead was funded by the National Health and Medical Research Council of Australia (ID310670, ID628903); the Cancer Institute NSW (12/RIG/1-17, 15/RIG/1-16); the Department of Gynaecological Oncology, Westmead Hospital; and acknowledges financial support from the SW-TCRC funded by the Cancer Institute NSW (15/TRC/1-01). AA-M is supported by the Guerrera Familly Cancer Scientist Award. We gratefully acknowledge the contributions of Catherine Kennedy, Jessica Boros, Yoke-Eng Chiew, the Westmead Hospital Department of Gynaecological Oncology, clinical collaborators and all the women who have consented to participate in research through the Westmead GynBiobank. The JGH Breast Biobank is supported by the RRCancer and the Quebec Breast Cancer Foundation. Figures created with BioRender.com. Support for title page creation and format was provided by AuthorArranger, a tool developed at the National Cancer Institute.

## Author Contributions

Conceptualization: QZ, RR. Methodology: ZC, ZN, EB, PH, QZ. Software: ZC, MD. Formal analysis: ZC, ZN, EB, AA, QZ. Investigation: ZC, ZN, EB, AA, QZ. Data generation and curation: all authors. Writing – original draft: ZC, EB, QZ. Writing – review and editing: all authors

## Supplementary Information

**Supplementary Table 1.**
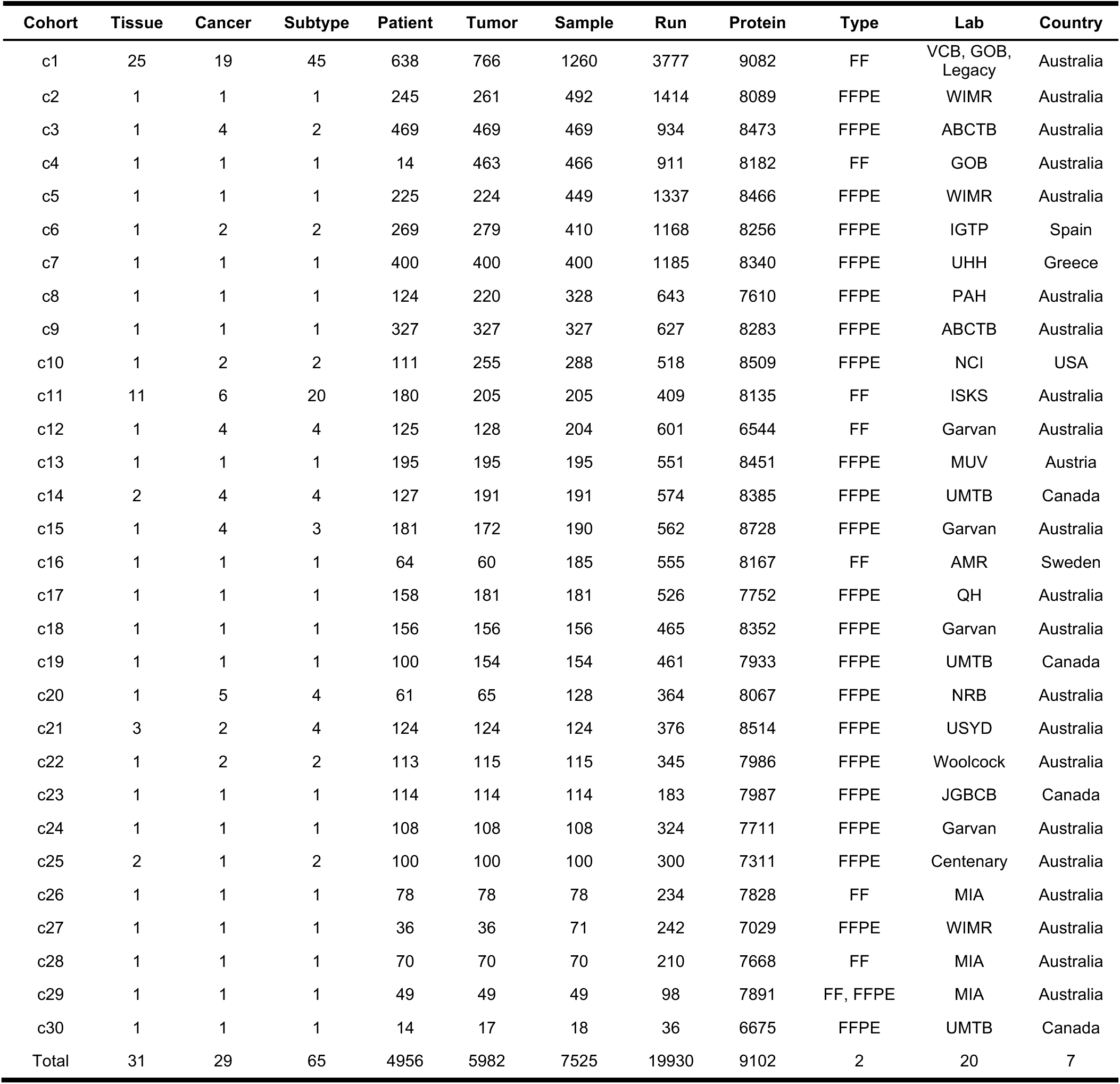
Cohort details of ProCan Compendium. Tissue refers to tissue of origin. Run represents the number of DIA-MS runs. Protein means the number of quantified proteins.

**Supplementary Table 2.**
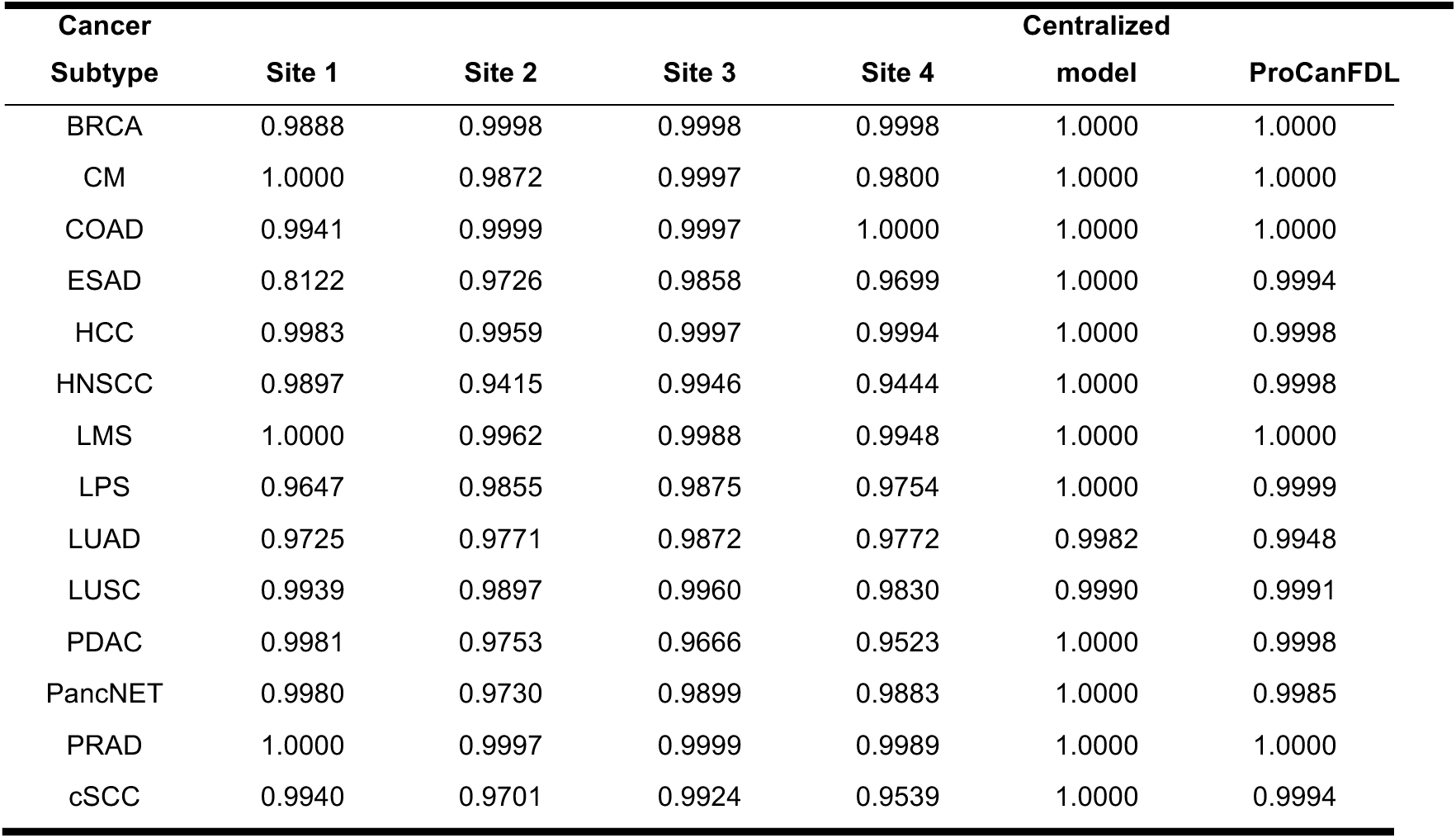
Per class AUROC for models using ProCan Compendium.

**Supplementary Table 3.** Cohort details of external proteomic datasets.

| Cohort | Subtypes | Patients | Proteins | Labs | Country |
| --- | --- | --- | --- | --- | --- |
| DIA-1 | 1 | 15 | 4238 | University of Witwatersrand | South Africa |
| DIA-2 | 1 | 40 | 4238 | IMIBIC | Spain |
| CPTAC-BRCA | 1 | 122 | 7748 | CPTAC | USA |
| CPTAC-PDAC | 1 | 105 | 7741 | CPTAC | USA |
| CPTAC-LUAD | 1 | 106 | 7871 | CPTAC | USA |
| CPTAC-HNSCC | 1 | 108 | 7751 | CPTAC | USA |
| CPTAC-COAD | 1 | 97 | 7086 | CPTAC | USA |
| CPTAC-LSCC | 1 | 108 | 7940 | CPTAC | USA |
| CPTAC-OV | 1 | 83 | 7390 | CPTAC | USA |
| CPTAC-CCRCC | 1 | 103 | 7589 | CPTAC | USA |
| Total | 10 | 887 | 8594 | 3 | 3 |

**Supplementary Table 4.**
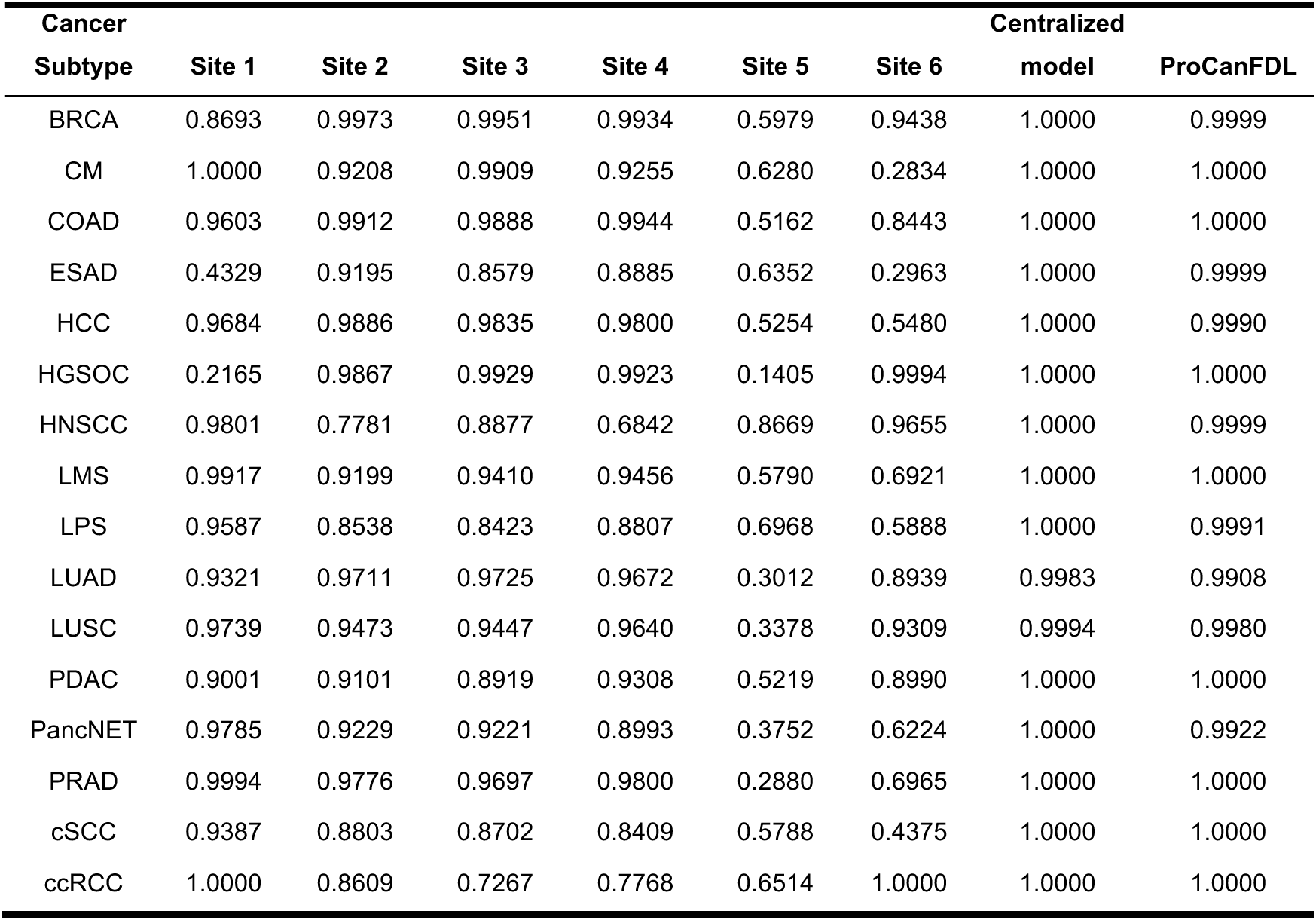
Per class AUROC using ProCan Compendium and external datasets.

**Supplementary Figure 1.**
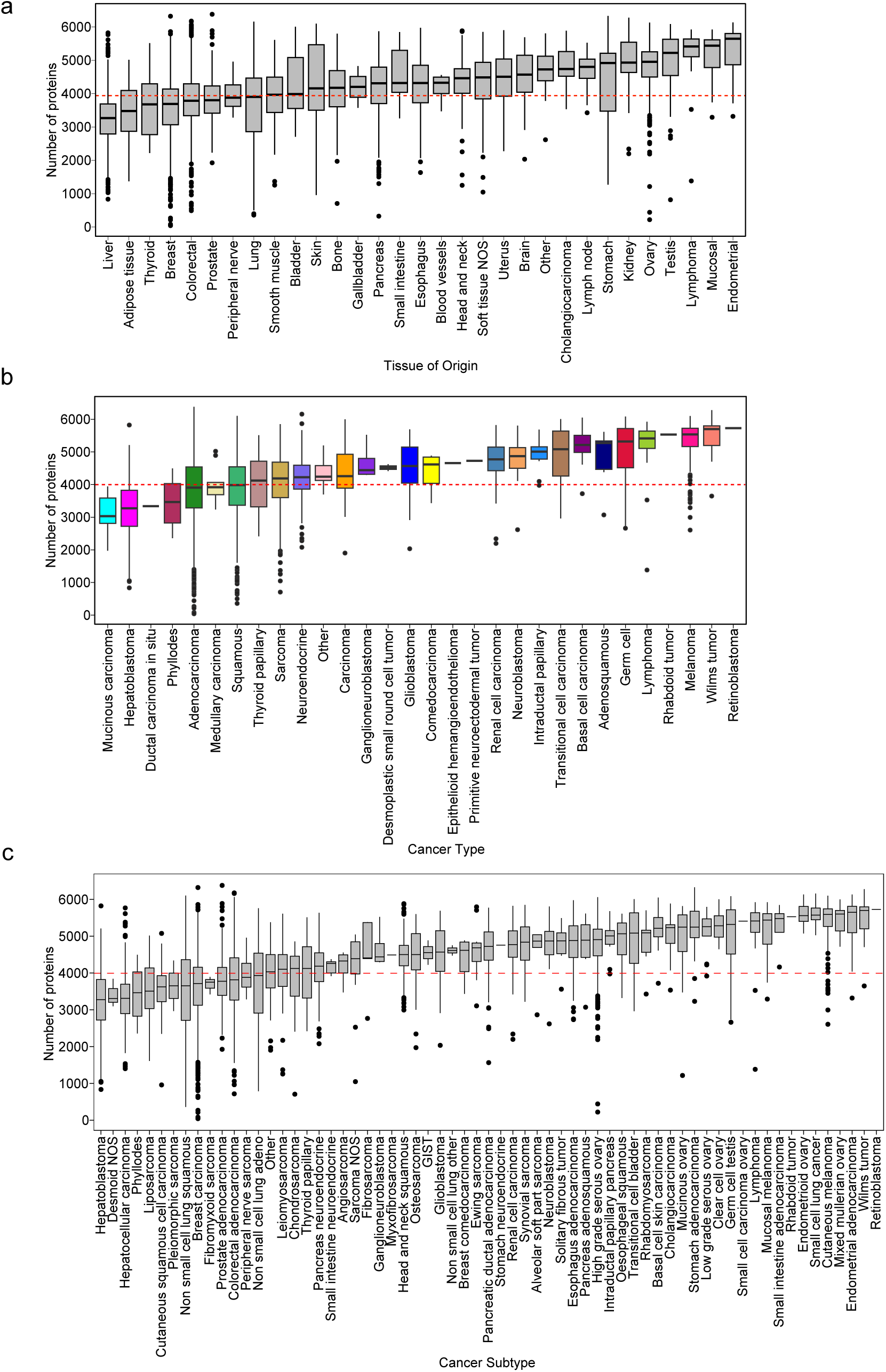
Overview of ProCan Compendium. **a)** Number of proteins quantified per sample, grouped by tissue of origins. **b)** grouped by cancer types. **c)** grouped by cancer subtypes. NOS, not otherwise specified. The red line in each of the three plots shows the median protein count per sample.

**Supplementary Figure 2.**
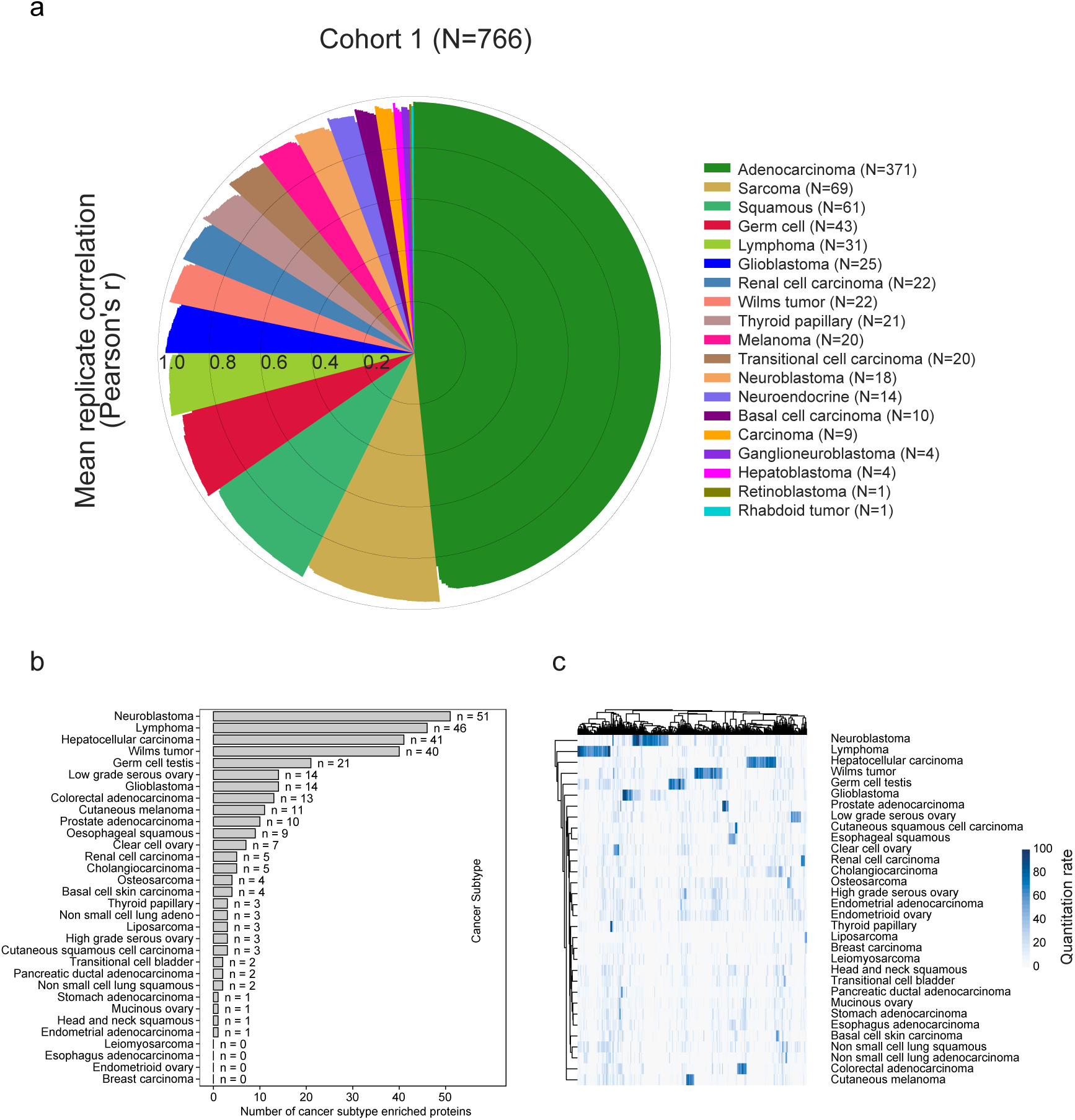
Overview of Cohort 1. **a)** Mean Pearson’s correlation for replicates of each sample, colored by cancer types. **b**) Bar plot displaying the number of cancer subtype-enriched proteins for each cancer subtype. Enriched proteins are defined as those present in >50% of samples from no more than one cancer subtype and in 35% or fewer samples from all other subtypes, with each subtype having at least ten samples. **c**) Heatmap with hierarchical clustering of cancer subtype-enriched proteins, with the quantification rate indicated by the color gradient. Rows represent cancer subtypes, and columns represent proteins. The quantification rate shows the percentage of samples in which the protein is quantified for each cancer subtype.

**Supplementary Figure 3.**
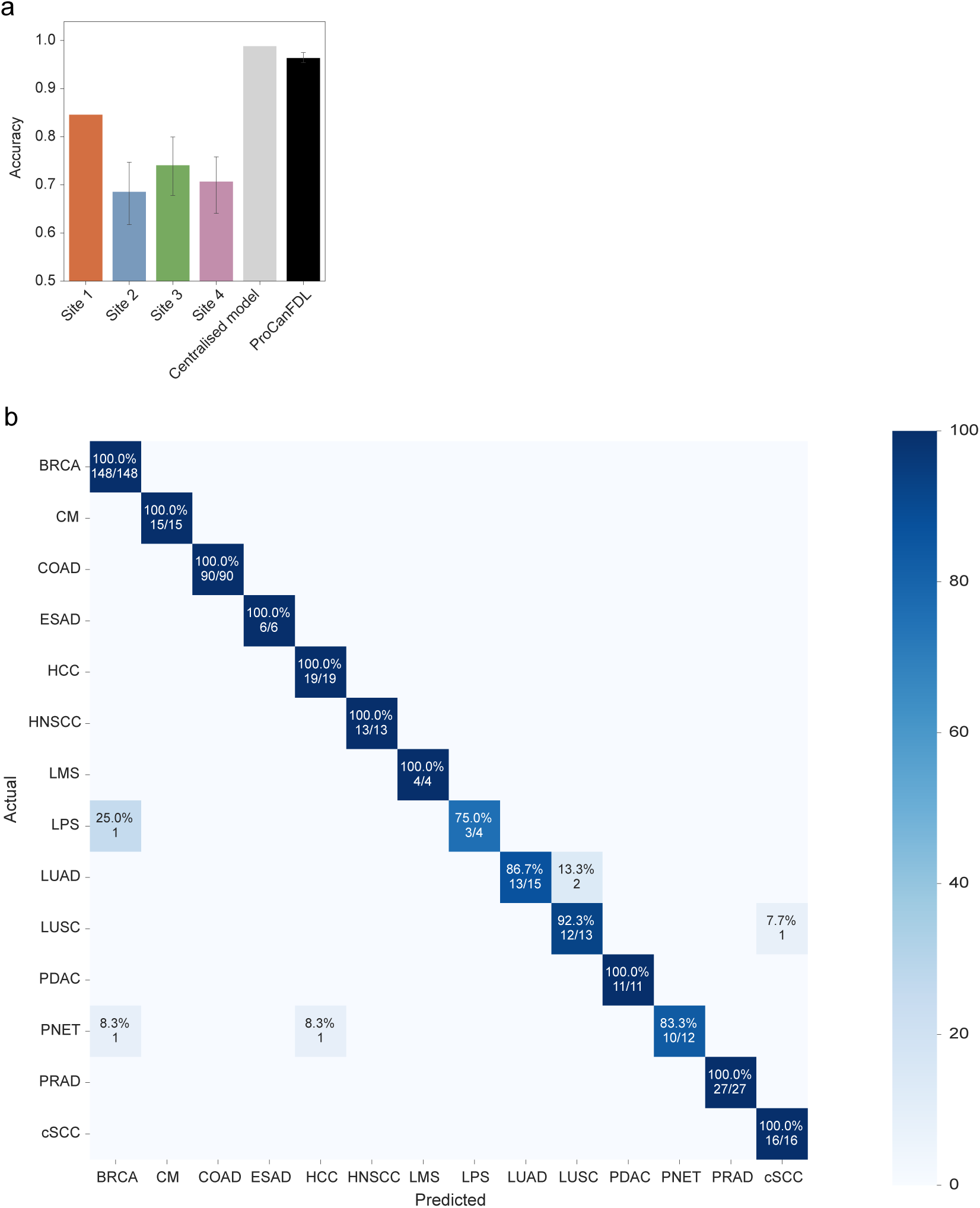
ProCanFDL of ProCan Compendium. **a)** Benchmark by accuracy for different models. Error bars for Sites 2-4 and ProCanFDL represent 95% CI estimated from ten experiments. **b**) A confusion matrix shows the number of correct predictions, with the total number of cases annotated using blue shading. Percentage represents the sensitivity for each cancer subtype.

**Supplementary Figure 4.**
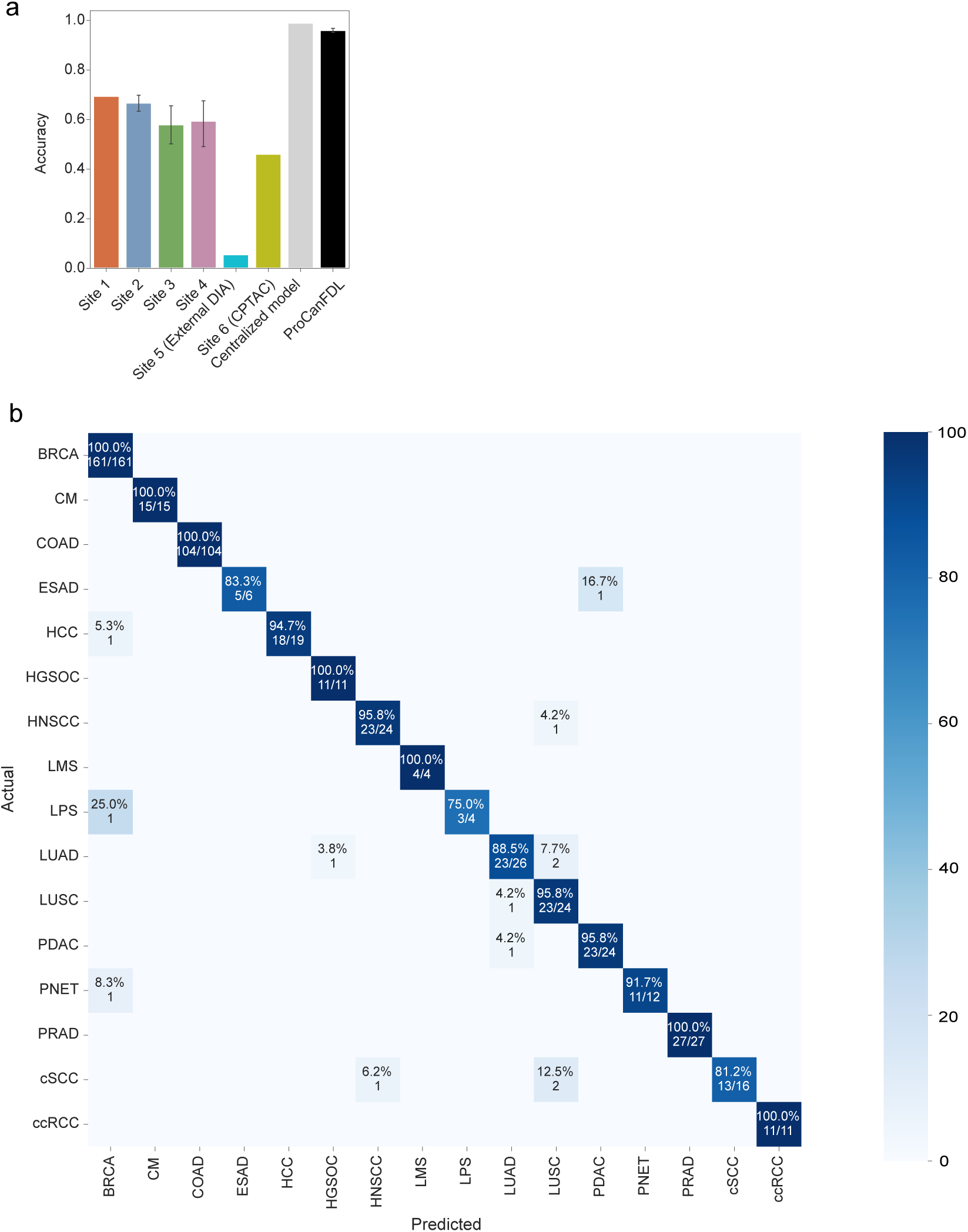
Effects of adding external datasets. **a)** Accuracy of different models. Error bars represent 95% CI. **b)** A confusion matrix shows the number of correct predictions with the total number of cases annotated using blue shading. Percentage represents the sensitivity for each cancer subtype.

**Supplementary Figure 5.**
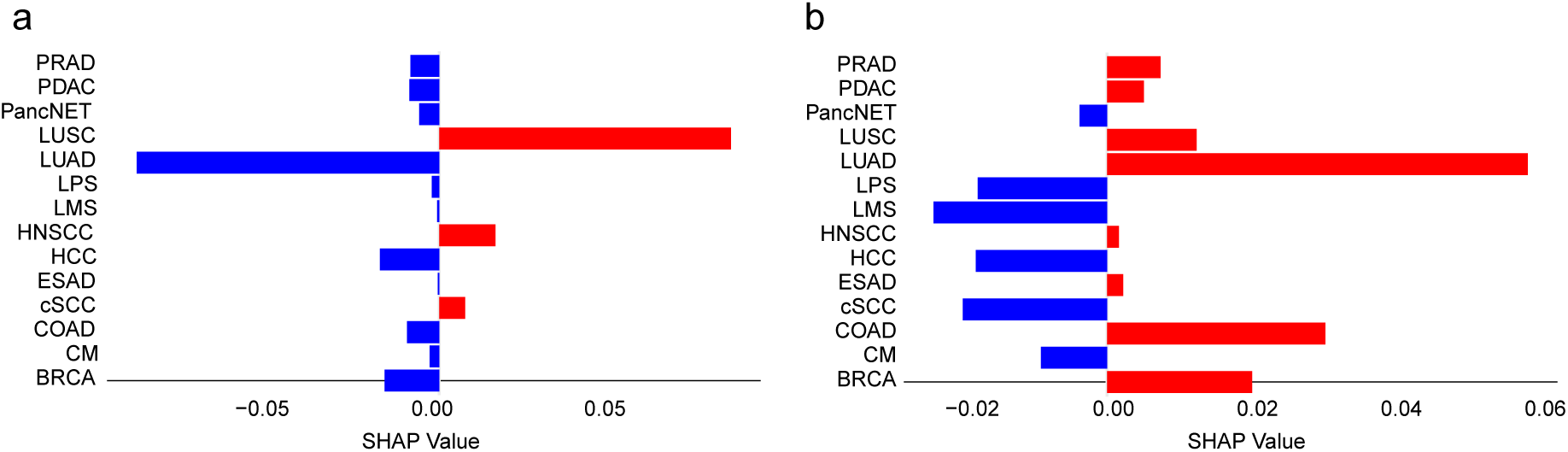
Feature importance for selected proteins with utility at distinguishing cancer subtypes. Features contributing positively to class prediction shown on the right-hand side and features contributing negatively on the left-hand side. **a)** SHAP values for desmoglein 3 (DSG3) across 14 cancer subtypes **b)** SHAP values for Anterior gradient 2, protein disulfide isomerase family member (AGR2) across 14 cancer subtypes.

